# The intensity of the immune response to LPS and *E. coli* regulates the induction of preterm labor in Rhesus Macaques

**DOI:** 10.1101/2021.01.07.425700

**Authors:** Monica Cappelletti, Pietro Presicce, Feyaing Ma, Paranthaman Senthamaraikannan, Lisa A. Miller, Matteo Pellegrini, Alan H. Jobe, Senad Divanovic, Sing Sing Way, Claire A. Chougnet, Suhas G. Kallapur

**Affiliations:** Divisions of Neonatology and Developmental Biology, David Geffen School of Medicine at the University of California Los Angeles, CA, USA; Department of Molecular, Cell and Developmental Biology Medicine at the University of California Los Angeles, CA, USA; Institute for Quantitative and Computational Biosciences – Collaboratory at the University of California Los Angeles, CA, USA; California National Primate Research Center UCD, CA; Division of Neonatology/Pulmonary Biology, Cincinnati Children’s Hospital Research Foundation, and the University of Cincinnati College of Medicine, Cincinnati, OH, USA; Immunobiology, Cincinnati Children’s Hospital Research Foundation, and the University of Cincinnati College of Medicine, Cincinnati, OH, USA; Infectious Diseases, Cincinnati Children’s Hospital Research Foundation, and the University of Cincinnati College of Medicine, Cincinnati, OH, USA; California National Primate Research Center, University of California Davis, Davis, CA, Department of Anatomy, Physiology, and Cell Biology, School of Veterinary Medicine, University of California Davis, CA, USA

**Author notes:** **Correspondence:** Suhas G. Kallapur, MD, Chief, Divisions of Neonatology and Developmental Biology, Professor of Pediatrics, David Geffen School of Medicine at UCLA, Mattel Children’s Hospital UCLA, 10833 Le Conte Avenue, Room B2-375 MDCC, Los Angeles, CA 90095, Email id.

**Keywords:** chorioamnionitis, neutrophils, inflammation, innate immunity, reproductive immunology

## Abstract

Intrauterine infection/inflammation (IUI) is a major contributor to preterm labor (PTL). However, IUI does not invariably cause PTL. We hypothesized that quantitative and qualitative differences in immune response exist in subjects with or without PTL. To define the triggers for PTL, we developed Rhesus macaque models of IUI driven by lipopolysaccharyde (LPS) or live *E. coli*. PTL did not occur in LPS challenged Rhesus macaque while *E. coli* infected animals frequently delivered preterm. Although LPS and live *E. coli* both caused immune cell infiltration, *E. coli* infected animals showed higher levels of inflammatory mediators, particularly IL6 and prostaglandins, in the chorioamnion decidua and amniotic fluid. Neutrophil infiltration in the chorion was a common feature to both LPS and *E. coli*. However, neutrophilic infiltration and *IL6* and *PTGS2* expression in the amnion was specifically induced by live *E. coli*. RNASeq analysis of fetal membranes revealed that specific pathways involved in augmentation of inflammation including type I interferon response, chemotaxis, sumoylation and iron homeostasis were upregulated in the *E. coli* group compared to the LPS group. Our data suggest that intensity of the host immune response to IUI may determine susceptibility to PTL.

## INTRODUCTION

Intrauterine infection/inflammation (IUI), most commonly originates in the lower genitourinary tract and ascends to the uterine cavity including the chorio-decidual space and the amniotic cavity. About 40% of preterm labor (PTL) cases are associated with IUI (1). Importantly, the causal link between IUI and PTL is well-established (2). In a prospective preterm labor biomarker study, only 25% of women with increased amniotic fluid (AF) IL-6 had detection of microorganisms in the AF (3). Thus, both intrauterine infection and inflammation can cause PTL. Currently, the rate of prematurity remains stubbornly high at about 10% of all US births (4), causing 75% of perinatal mortality and 50% of the long-term morbidity (1). Apart from maternal morbidity, IUI causes fetal inflammation and increases the risk for fetal and newborn brain, gut, and lung injury in both clinical studies and in animal models (reviewed in (5)). Although IUI is a frequent cause of PTL, about 25-45% of pregnancies at risk for PTL (including cases with presumed IUI) do not delivery preterm within 14d of presentation (3, 6–8). Thus, PTL is not an invariable consequence of IUI.

What factors during IUI lead to crossing the threshold of PTL are not known. Human studies of PTL have implicated proinflammatory agents such as interleukin-1 (IL-1), interleukin-6 (IL-6), interleukin-8 (IL-8) and tumor necrosis factor-α (TNFα), Prostaglandins, Metalloproteinases, and the alarmin HMGB1 as potential etiologic factors (2, 3, 9, 10). However, large unbiased proteomic or genome wide association studies have not yielded clinical useful biomarkers of PTL (11, 12). A caveat with transcriptomic studies in both humans and animal models of PTL is that the controls often have no or very little inflammation (13, 14). Pathways leading to IUI tend to confound the analyses of PTL.

We previously reported that intraamniotic injection of LPS (from *E. coli*) or IL1β in pregnant Rhesus macaques caused IUI but not PTL (15–17). We now report that intraamniotic (IA) injection of live *E. coli* causes IUI and PTL. Importantly, live *E. coli*-induced PTL was not rescued by antibiotics, recapitulating the lack of efficacy of prenatal antibiotics in reducing PTL in humans (18, 19).

Thus, our two models of LPS-induced IUI, and live *E. coli*-induced PTL with IUI offer unique opportunities to unravel the IUI specific pathways responsible for induction of PTL. We tested the hypothesis that quantitative and qualitative thresholds in immune response exist in subjects with or without PTL.

## RESULTS

### New Rhesus model of IUI with PTL: Intraamniotic *E. coli* injection

*E. coli* was used for the studies since this organism is a prototypic invasive perinatal pathogen and a major cause of early neonatal sepsis resulting from maternal IUI (20). In our new model of IUI, pregnant Rhesus macaque were inoculated intraamniotically (IA) with live *E. coli* (1 x 10^6^ CFU). PTL was monitored for 48h after LPS or live *E. coli* after which surgical delivery was performed. In contrast to IA LPS causing no PTL within 48h, infection with IA *E. coli* caused PTL in 5/5 dams within 48h of injection (**Table 1**). To confirm lack of PTL in the IA LPS group, we extended the period of observation to 5d in another set and confirmed no PTL in 0 out of 9 animals (**Table 1**). Furthermore, IA *E. coli* caused maternal bacteremia in 3/5 dams and fetal bacteremia in 100% of pregnancies (**Table 2**).

**Table 1:**
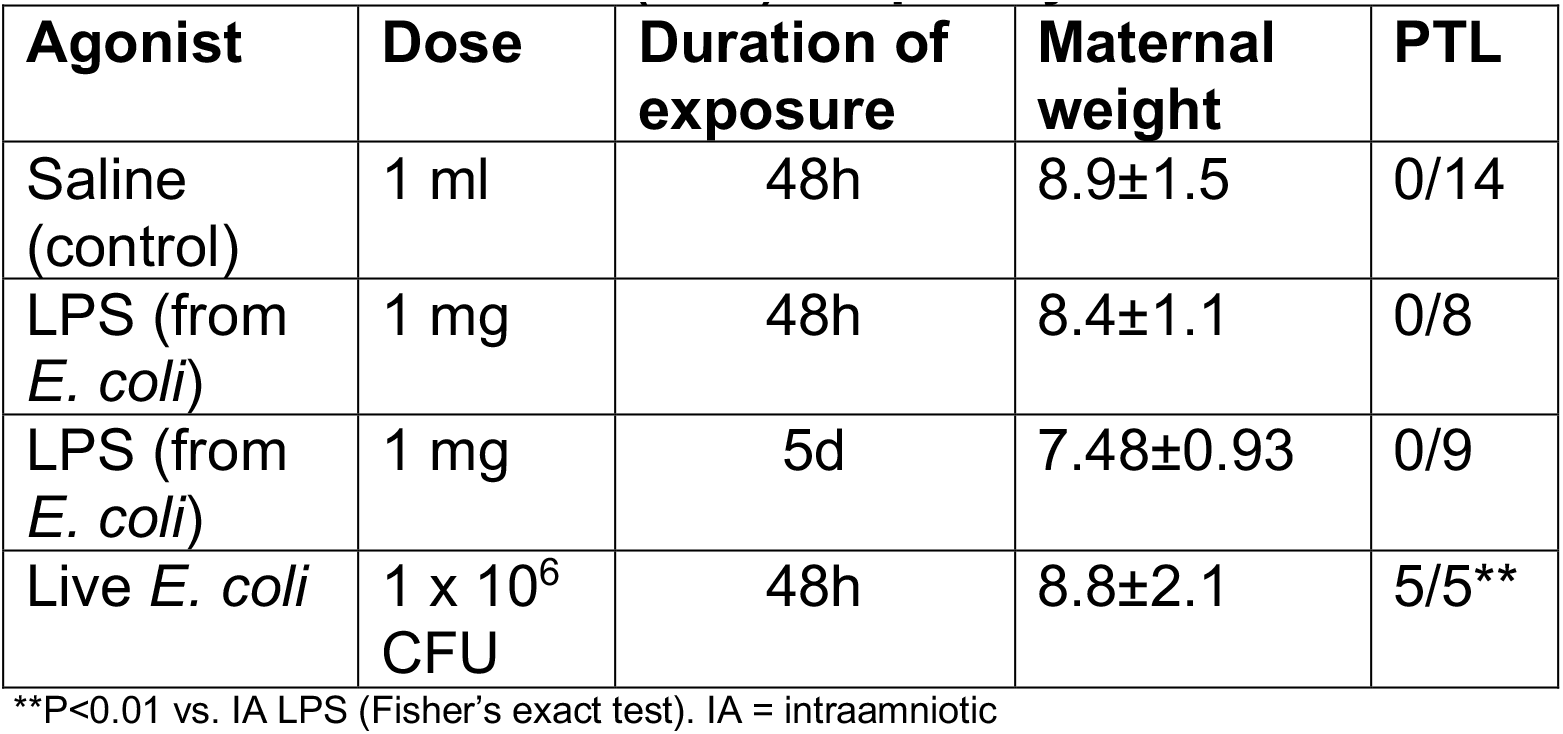
Preterm labor (PTL) frequency after IUI in Rhesus

| **Agonist** | **Dose** | **Duration of exposure** | **Maternal weight** | **PTL** |
| --- | --- | --- | --- | --- |
| Saline (control) | 1 ml | 48h | 8.9±1.5 | 0/14 |
| LPS (from *E. coli*) | 1 mg | 48h | 8.4±1.1 | 0/8 |
| LPS (from *E. coli*) | 1 mg | 5d | 7.48±0.93 | 0/9 |
| Live *E. coli* | 1 x 10<sup>6</sup> CFU | 48h | 8.8±2.1 | 5/5** |

\*\*P<0.01 vs. IA LPS (Fisher's exact test). IA = intraamniotic

**Table 2:** *E. coli* invasion in tissue compartments

| **Agonist** | **AF culture + (at delivery)** | **CD culture +** | **Maternal blood culture +** | **Fetal blood or liver culture +** |
| --- | --- | --- | --- | --- |
| Live *E. coli* | 3/3 | NA | 3/5 | 5/5 |

AF= Amniotic fluid, CD=chorio decidua

### LPS and *E. coli* induce similar immune infiltration in the chorioamnion decidua

To characterize IUI, we analyzed immune cell infiltration in the chorioamnion-decidua in different Rhesus models of IUI. Neutrophils were not detected in the controls but readily detected in the chorio-decidua after LPS or *E. coli* exposure (**Figure 1A**). Additionally, *E. coli* induced neutrophil infiltration in the amnion (**Figure 1A**). Flow cytometry demonstrated that neutrophils were the most abundant immune cells after LPS and *E. coli* exposures, in terms of both frequency (**Figure 1B**) and absolute counts (**Supplementary Figure 1**) (Gating strategy is shown in **Supplementary Figure 2**). Neutrophil frequency and counts were higher after *E. coli* compared to LPS exposure. Macrophages and NK cell frequency decreased after either LPS or *E. coli* exposure but absolute counts were similar to controls. T-cell frequency was similar among all groups (**Figure 1B**) but the absolute counts were higher in the IA *E. coli* group compared to saline controls (**Supplementary Figure 1**). NKT cells and B cell frequency or absolute counts did not significantly change in the LPS and *E. coli* groups (**Figure 1B**). Overall, both models induced similar immune infiltration with a higher neutrophil frequency after IA *E. coli* compared to IA LPS exposure.

**Figure 1.**
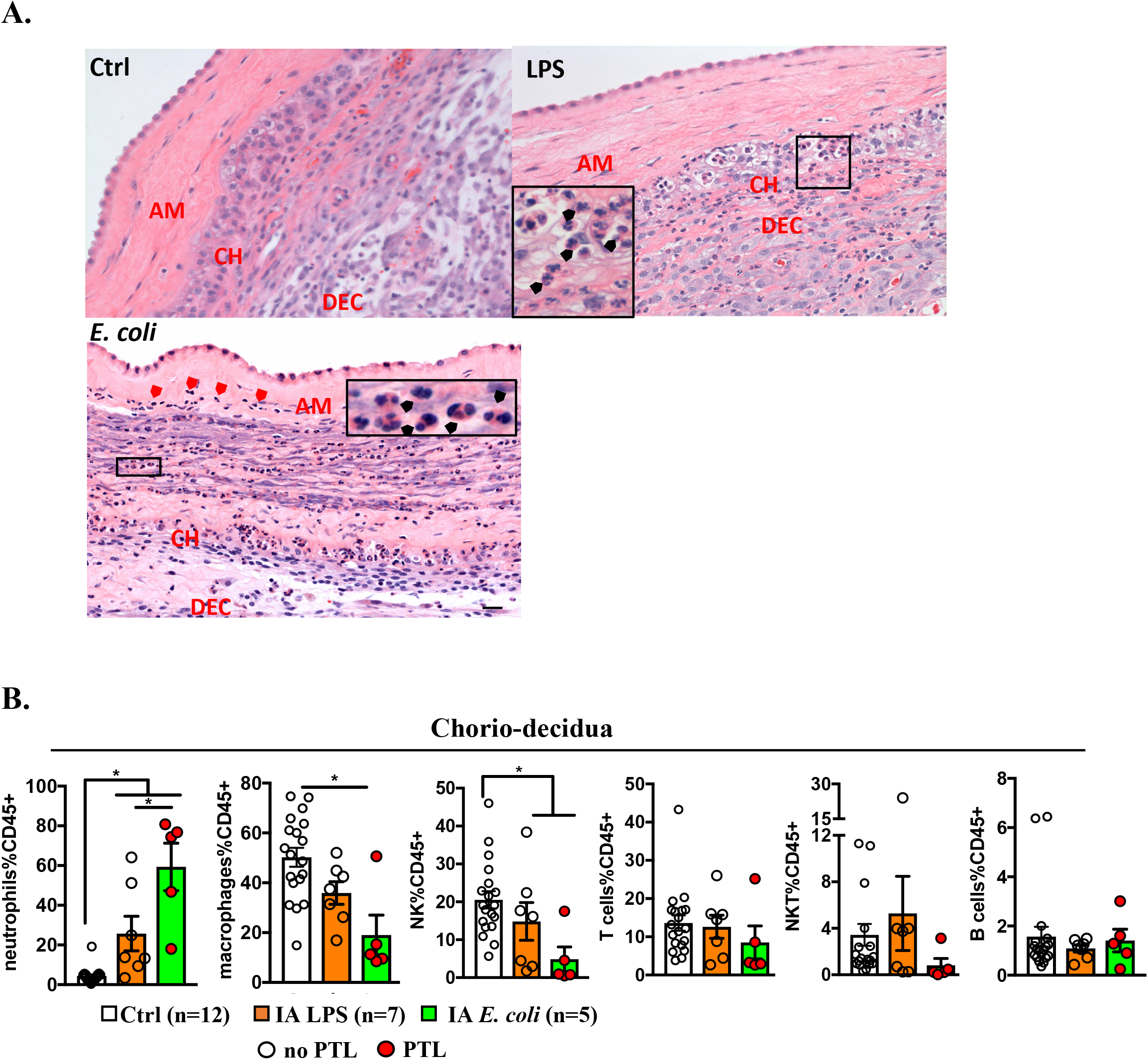
Increased neutrophil infiltration in chorio-decidua upon IA LPS or *E. coli* exposure. (**A**) Representative Hematoxylin & Eosin (H&E) staining showing neutrophil infiltration in fetal membranes (chorion, amnion, and decidua) after LPS or live *E. coli* exposure but not in controls. Note the LPS induced accumulation of neutrophils at the chorio-decidua junction (black arrows) but not in the amnion. IA *E. coli* injection induced neutrophil infiltration in the amnion (red arrows) in addition to the chorio-decidua (black arrows). AM (amnion); CH (chorion); and DEC (decidua). (**B**) Chorio-decidua cell suspensions were analyzed by multiparameter flow cytometry and the different leukocyte populations were defined as indicated in Supplementary Figure 2. IA *E. coli* exposure increased significantly the frequency of chorio-decidua neutrophils, that represent the major immune cell population. A decrease in frequency of macrophages and NK cells with no changes for T cells and B cells compared to the control animals is observed. Frequency is expressed as percent of CD45+ cells. In the IA *E. coli* group (green bars), red circles denote animals with PTL while clear circles denote animals without PTL. Data are mean, SEM, *p<0.05 between comparators (Mann–Whitney U-test).

### *E. coli* induces higher proinflammatory cytokines and prostaglandins in the AF compared to LPS

We determined the level of inflammation induced in the different IUI models. In comparison to controls, all of the mediators tested increased after LPS or *E. coli* exposure in the chorioamnion decidua with the exception of *IL6*, which only increased in the *E. coli* group (**Figure 2A**). *IL1β, CCL2, IL6* and *PTGS2* expression significantly increased in the chorioamnion-decidua of *E. coli* group compared to LPS (**Figure 2A**). Endotoxin levels in the amniotic fluid were significantly higher after IA *E. coli* compared to IA LPS (**Figure 2B**). Given that neutrophils were the predominant immune cells in the inflamed chorioamnion-decidua, we next examined their levels in the amniotic fluid (AF). The numbers of AF neutrophils increased comparably in both *E. coli* and LPS groups (**Figure 2C**). We then compared the cytokine responses in the AF of *E. coli* vs LPS groups. Levels of AF IL-1β, TNFα, IL-6 were all significantly higher in animals inoculated with *E. coli* and LPS groups compared to controls, with a trend towards higher levels in the *E. coli* compared to LPS group (**Figure 2D**). Similarly, the prostaglandins PGE2 and PGF2α increased in both groups compared to controls but we observed 2-3 fold higher concentration in the AF from *E. coli* vs LPS animals (**Figure 2E**). In contrast to large increases of cytokines in the AF, the changes in maternal plasma were much more modest. *E. coli* and LPS exposure increased IL-6 and CCL2 levels in the maternal plasma but did not change IL8 or TNFα levels (**Supplementary Figure 3A**). The fetal plasma cytokines increased slightly but significantly after LPS. Although we only had 2 samples available, *E. coli* exposure also increased fetal plasma cytokines (**Supplementary Figure 3B**).

**Figure 2.**
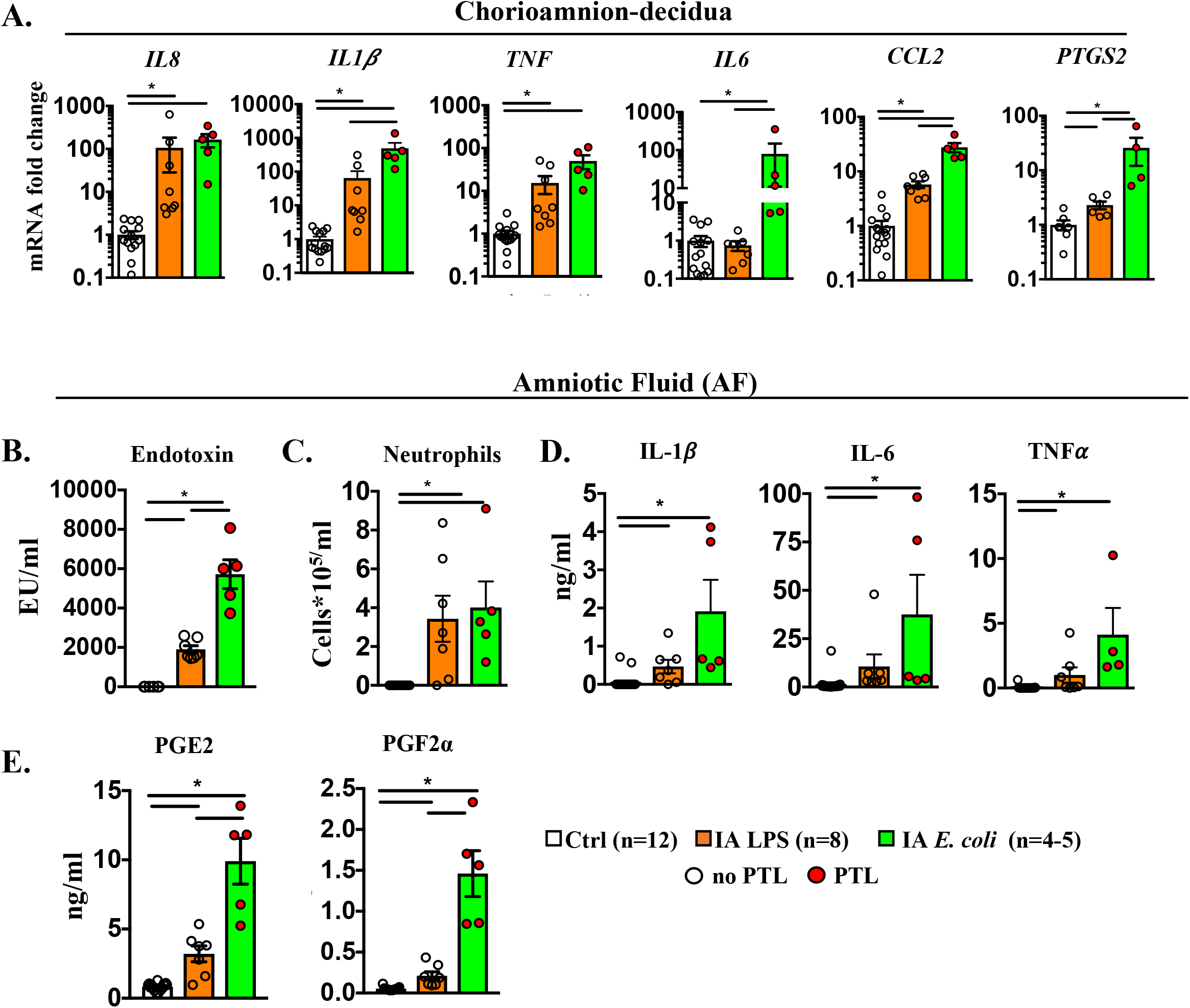
Higher inflammation in the Chorioamnion-decidua and in the Amniotic Fluid by live *E. coli* compared to LPS. (**A**) Chorio-amnion decidua inflammatory cytokine mRNAs in different Rhesus IUI models were measured by qPCR. Average mRNA values are fold increases over the average value for the control group after internally normalizing to the housekeeping 18S RNA. Amniotic fluid levels of (**B**) endotoxin levels measured by limulus lysate assay, (**C**) neutrophil frequency measured by cell and differential counts on cytospins (**D**) inflammatory cytokines, and (**E**) prostaglandins. Compared to IA LPS-exposure, IA E. coli-exposure induced a higher expression of IL1b, IL6, CCL2 and PTGS2 mRNAs in the chorioamnion-decidua, higher AF endotoxin levels, and higher prostaglandin levels in the AF. Data are mean, SEM, *p<0.05 between comparators. In the E. coli group (green bar), red circles denote animals with PTL, clear circles denote animals without PTL. *p < 0.05 (Mann–Whitney U-test).

Differential upregulation of mediators in the *E. coli* vs. LPS groups could be due to time dependent effects. We previously reported that IA LPS induction of intrauterine inflammation is higher at 16h compared to 48h (21). We therefore compared key genes differentially expressed in the *E. coli* group with LPS exposure of 16h (using samples archived from our previous study (21)). Similar to the results for the IA LPS 48h group, *E. coli* induction of chorioamnion-decidua expression of *IL6* and *CCL2* were higher compared to controls and AF prostaglandin levels were higher compared to IA LPS 16h (**Supplementary Figure 4A-B**). These results suggest that higher induction of key mediators of IUI after IA *E. coli* compared to LPS may not be explained by temporal trajectories of gene expression.

### *E. coli* drives higher levels of inflammation in the amnion

Because *E. coli* but not LPS induced neutrophil infiltration in the amnion (**Figure 1A**), we analyzed the anatomic locations of inflammatory response. We surgically separated the amnion from the chorio-decidua. In comparison to controls, all of the mediators tested increased after LPS or *E. coli* exposure in the amnion with the exception of *IL6* which did not increase after LPS exposure (**Figure 3**). In comparison to the LPS group, *E. coli* increased amnion expression of *IL1β, IL8, TNFa, COX2, CCL2, IL6* and *PTGS2* mRNA levels. Together, these data show that inflammation in the amnion is significantly more pronounced after *E. coli* compared to LPS exposure.

**Figure 3.**
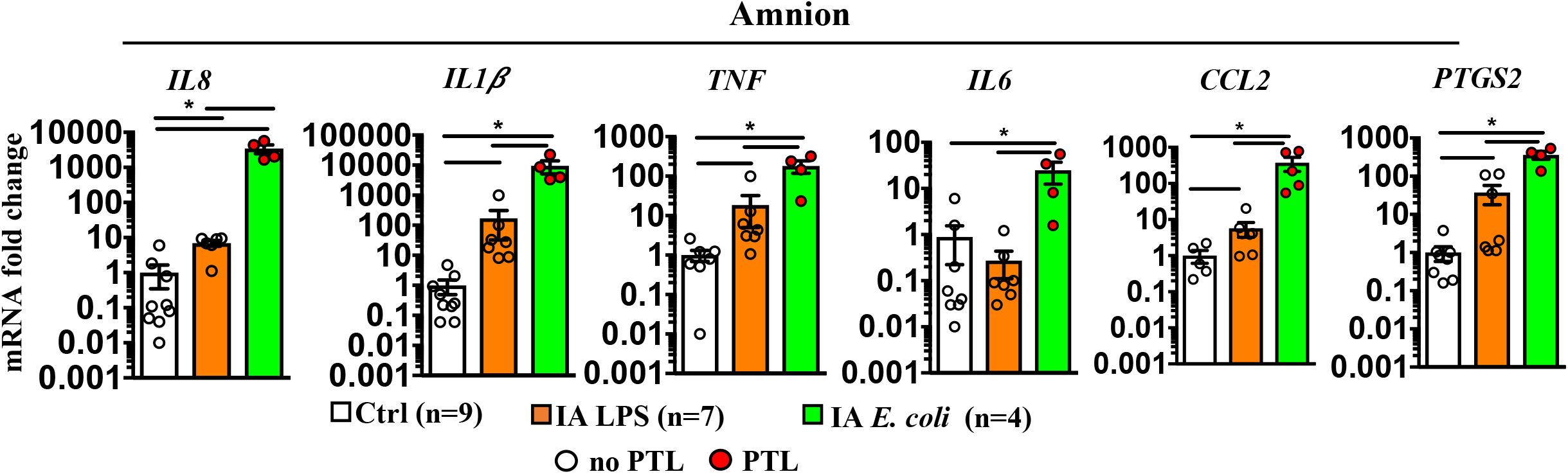
Higher inflammation in the Amnion by live *E. coli* compared to LPS. Inflammatory cytokine mRNAs in different Rhesus IUI models were measured in surgically isolated amnion tissue by qPCR. Average mRNA values are fold increases over the average value for the control group after internally normalizing to the housekeeping 18S RNA. Data are mean, SEM, *p<0.05 between comparators. In the E. coli group (green bar), red circles denote animals with PTL, clear circles denote animals without PTL. *p < 0.05 (Mann–Whitney U-test).

### Antibiotic treatment does not decrease *E. coli*-driven inflammation and preterm birth

To control bacteremia and thus simulate clinical situations, we added antibiotic treatment in pregnant Rhesus infected with *E. coli*. Cefazolin + enrofloxacin starting 24h after IA *E. coli* effectively eradicated maternal bacteremia in 7/8 subjects. In 1 subject, there was a persistent amniotic and fetal bacteremia. Despite the clearance of organisms in the antibiotics group, PTL was observed in 6/8 subjects by day 3. To understand if anti-microbial therapy also reduced inflammation, we compared select mediators in different compartments between IA *E. coli* and IA *E. coli* + antibiotics (Abx) groups. The tested mediators were comparable between the two groups: neutrophil frequency and *IL6* expression in the chorioamnion-decidua (**Figure 4A**), I*L6* expression in the amnion (**Figure 4B**) and IL-6 and PGE2 levels in the amniotic fluid (**Figure 4C**) were similar in the 2 groups.

**Figure 4.**
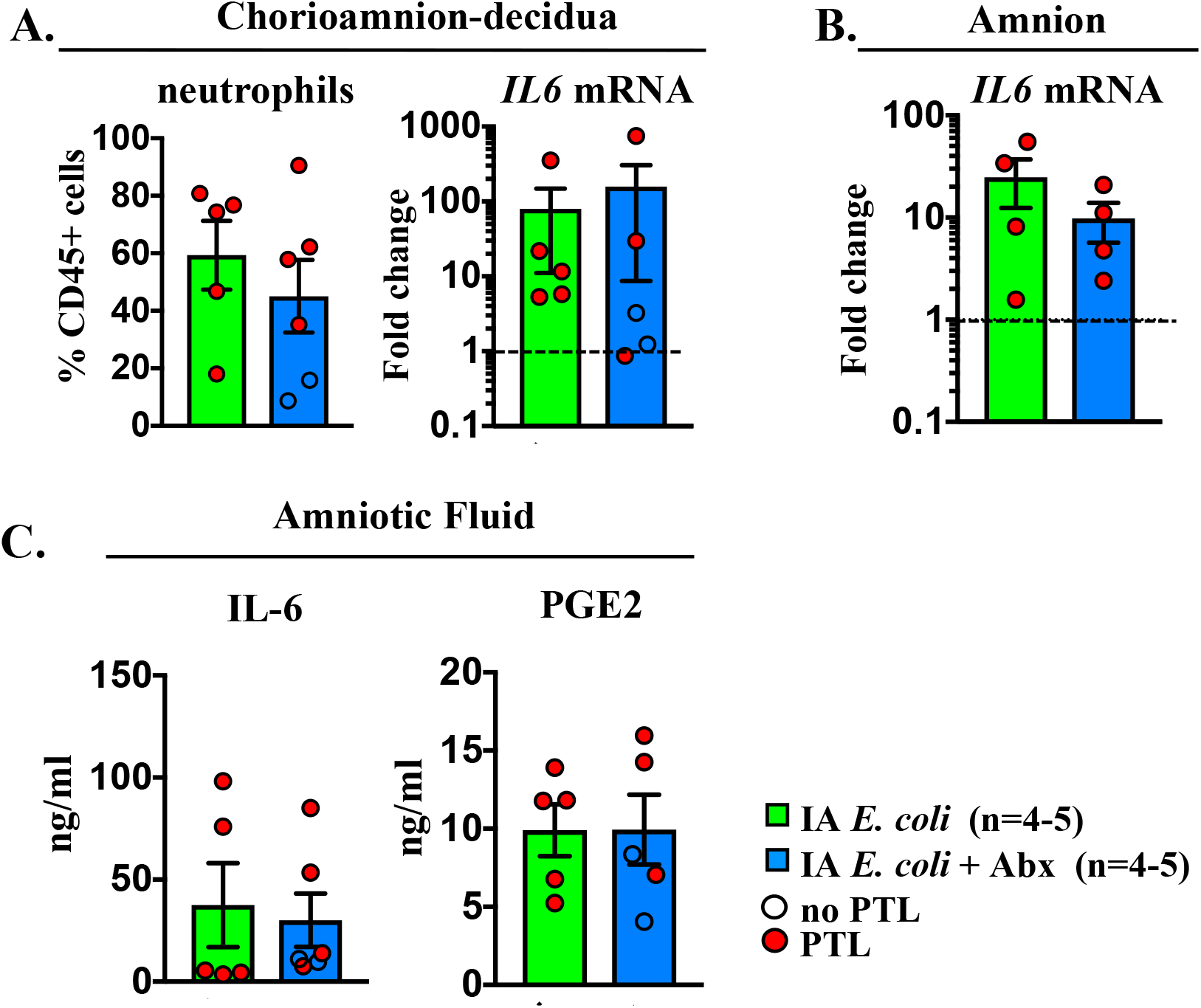
Antibiotic treatment (Abx) did not decrease *E. coli* induced inflammation. The comparison groups are IA E. coli without Abx (green bar) and IA *E.coli* with Abx (blue bar). Red circles denote animals with PTL, clear circles denote animals without PTL. Values were expressed as fold change value for the IA saline control group. Comparison of (**A**) chorio-amnion decidua neutrophil frequency and IL6 mRNA, (**B**) amnion IL6 mRNA, and (**C**) amniotic fluid IL6 and prostaglandin E2. Dashed line representing the mean value of ctrl animals. *p < 0.05 (Mann–Whitney U-test).

### Anatomical localization *of IL6* and *PTGS2* genes in the fetal membranes

Since *IL6* and prostaglandins were the key differentially expressed genes between *E. coli* +/- Abx and LPS groups, we colocalized IL6 or PTGS2 with MPO, a marker of activated neutrophils. We used dual RNAscope^®^ fluorescence *in situ* hybridization to visualize the *IL6* or *PTGS2* mRNA expression coupled with immunofluorescence to detect MPO^+^ cells. Sample from *E. coli* +/- Abx were combined because they did not show any significant differences (**Figure 4**). MPO^+^ cells infiltrated the chorio-decidua tissue similarly in both LPS and *E. coli* +/- Abx groups (**Figure 5A-B**), confirming the previous H&E findings (**Figure 1A**). However, in the amnion, MPO^+^ cells were rarely detected in the LPS group but abundant in the *E. coli* +/- Abx group (**Figure 5A-B**) again confirming the H&E findings (**Figure 1A**). The majority (>80-90%) of PTGS2^+^ cells were also MPO^+^ both in the LPS and *E. coli* groups, respectively, suggesting that chorio-decidua neutrophils are a major source of prostaglandin production in the fetal membranes during IUI (**Figure 5B**). MPO^+^ cells in the amnion of the *E. coli* exposed animals also expressed PTGS2, with far fewer MPO^+^PTGS2^+^ cells in the amnion of the LPS group. In contrast to the chorio-decidua being a major source of PTGS2 expression, IL6^+^ cells were exclusively present in the amnion of the *E. coli* +/- Abx group (**Figure 5C**) consistent with our rtPCR data (**Figure 3**). Notably, MPO^+^ cells did not express IL6. Rather, IL6^+^ cells morphologically appeared to be amnion epithelial cells and the amnion mesenchymal cells. Taken together, these observations indicate that neutrophil recruitment and PTGS2 expression in the chorion is a common feature to both LPS and *E. coli* +/- Abx immune response. However, neutrophilic PTGS2 expression and IL6 expression in the amnion is differentially induced by *E. coli* infection.

**Figure 5.**
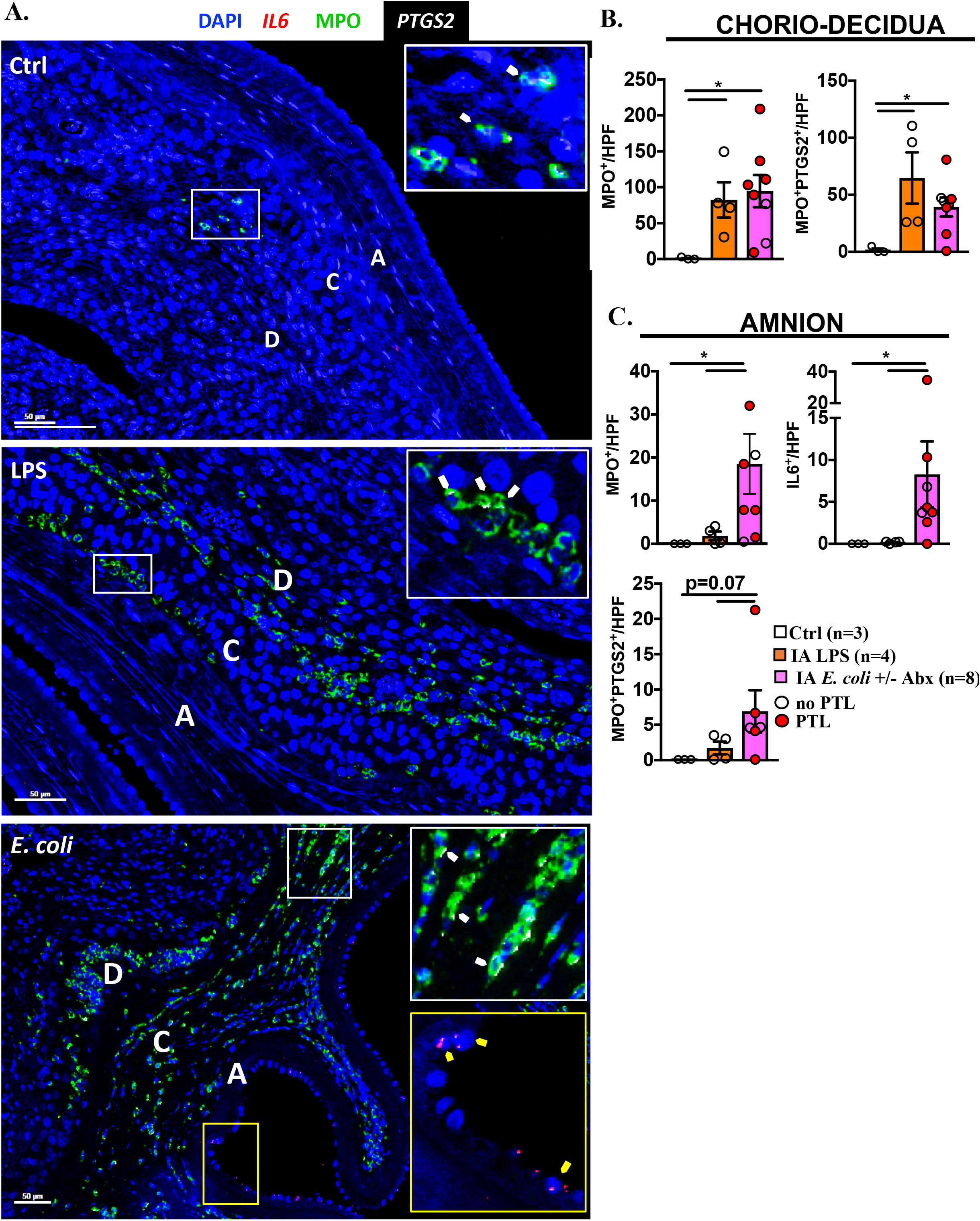
Cellular localization of IL6 and PTGS2 in the fetal membranes. (**A**) Representative multiplex fluorescence detection of IL6 and PTGS2 mRNA identified by RNA scope in situ hybridization and MPO co-localization by immunofluorescence. IL6 is shown in red, PTGS2 in white and MPO in green. DAPI indicates nuclear staining (blue) in all images. A=Amnion, C= Chorion, and D=Decidua. White arrows in the inset indicate co-localization of MPO (green) and PTGS2 (white) in the chorio-decidua. Yellow arrows in the inset indicate co-localization of MPO (green) and IL6 (red) in the amnion. Quantification of the cells expressing different markers in the (**B**) chorio-decidua, and (**C**) amnion. Average of 5 randomly selected high power field (HPF) fields were plotted as the representative value for the animal. Counts were performed in a blinded manner. Red circles denote animals with PTL. Clear circles denote animals without PTL. Data are mean, SEM, *p<0.05 between comparators (Mann-Whitney U-test).

### *E. coli* induces specific pathways in chorioamnion-decidua involved in exacerbation of inflammation

To gain transcriptomic insights associated with PTL, we compared RNA sequencing (RNA-seq) analysis using the chorioamnion-decidua from the different animal groups. For these studies, we combined RNA-seq analyses from *E. coli* (n=3) and *E. coli+Abx* (n=2) since both had similar inflammatory response (**Figure 4**). Unbiased principal component analysis (PCA) of global gene-expression profiles demonstrated distinct clustering of *E. coli* vs. LPS groups (**Figure 6A**). Interestingly, profiles for *E. coli* only and *E. coli* + Abx clustered together (**Figure 6A**), confirming our previous findings. Heat-maps of differentially expressed genes demonstrated distinct profiles in *E. coli* vs. LPS groups (**Figure 6B**). Importantly, the replicates within each group were similar, demonstrating validity of the findings (**Figure 6B**). Comparison of *E. coli* vs. LPS chorioamnion-decidua samples revealed 3,301 differentially expressed genes (DEGs) (fold change ≥ 2 and FDR adjusted p-value of 0.05) (**Figure 6C-D**). 1,415 genes were upregulated by *E. coli* compared to the LPS (**Figure 6D**). Gene-ontology category analysis revealed that the pathways upregulated by *E. coli* (representative genes) include: Positive regulation of chemotaxis (*CXCL3*), Type I Interferon α/β receptor signaling (*IRF7*), Global genome nucleotide-excision repair (*SUMO1*), and Iron homeostasis (*FHT1*) (**Figure 7A-B**). 1,102 genes were downregulated by *E. coli* compared to LPS. The major pathway downregulated by *E. coli* was post-translational modifications (e.g. ubiquitination and acetylation) (**Supplementary Figure 5A**). The unchanged genes were 784 and were involved in pathways regulating the mRNA processing, splicing and protein deacetylation (**Supplementary Figure 5B**).

**Figure 6.**
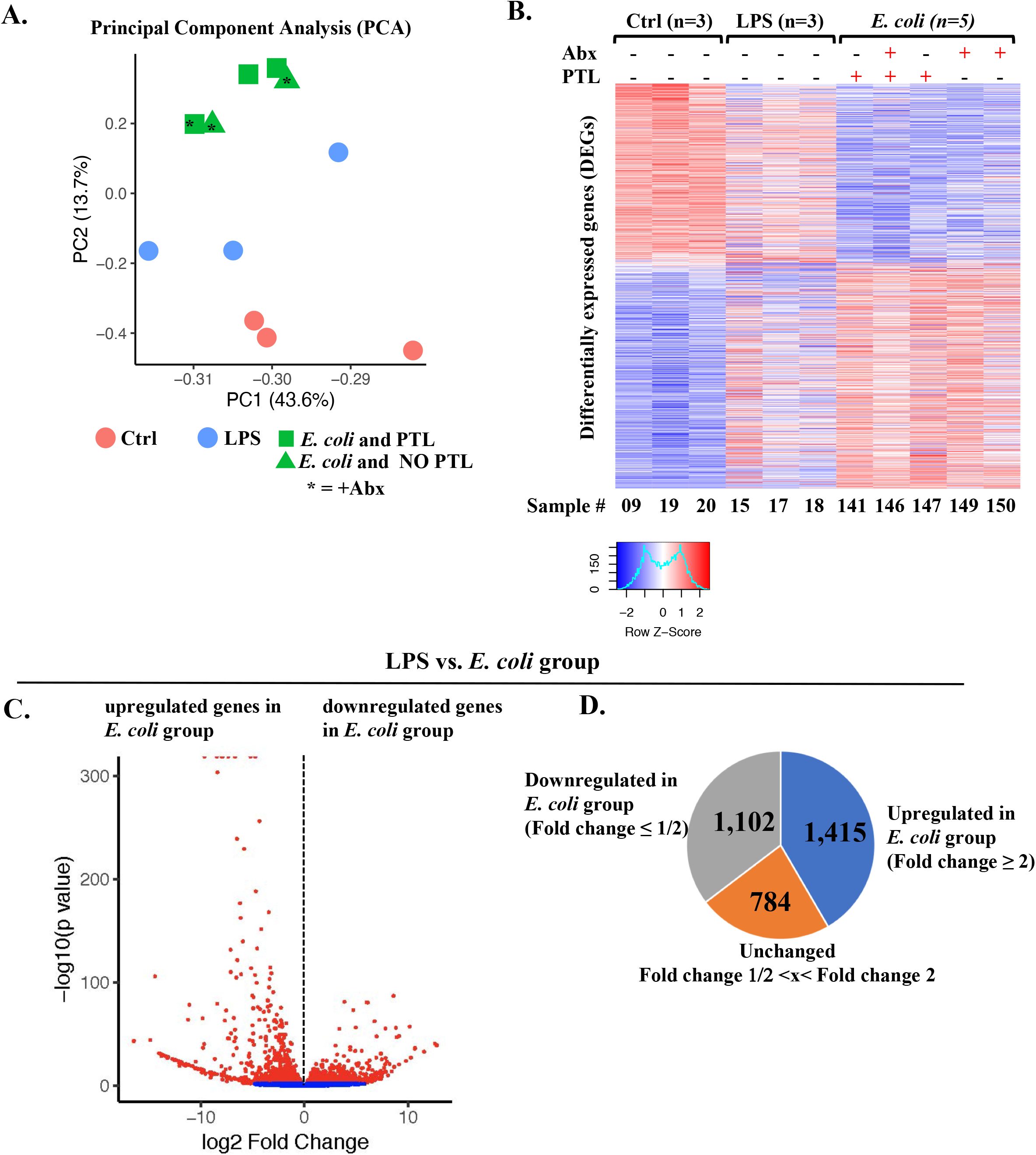
Comparative transcriptomics in chorioamnion-decidua between IA LPS- and *E. coli*-exposure. (**A**) Principal component analysis (PCA) of RNA-seq data from chorioamnion-decidua cells showing different clustering based on exposures. (**B**) Heatmap of genes that are differentially expressed in chorioamnion-decidua cells upon different treatments showing the minimal inter-animal variability within each group. (**C**) Volcano plot displaying differentially expressed genes from *E. coli* animals compared with LPS animals. Red dots indicate genes with DEGs > Fold Change 2 and FDR-adjusted p-value <0.05. (**D**) Pie chart displaying DEGs upregulated (Fold Change ≥ 2), unchanged (Fold Change 1/2<x< Fold Change 2) and downregulated in *E. coli* (Fold Change ≤ 1/2) compared to LPS.

**Figure 7.**
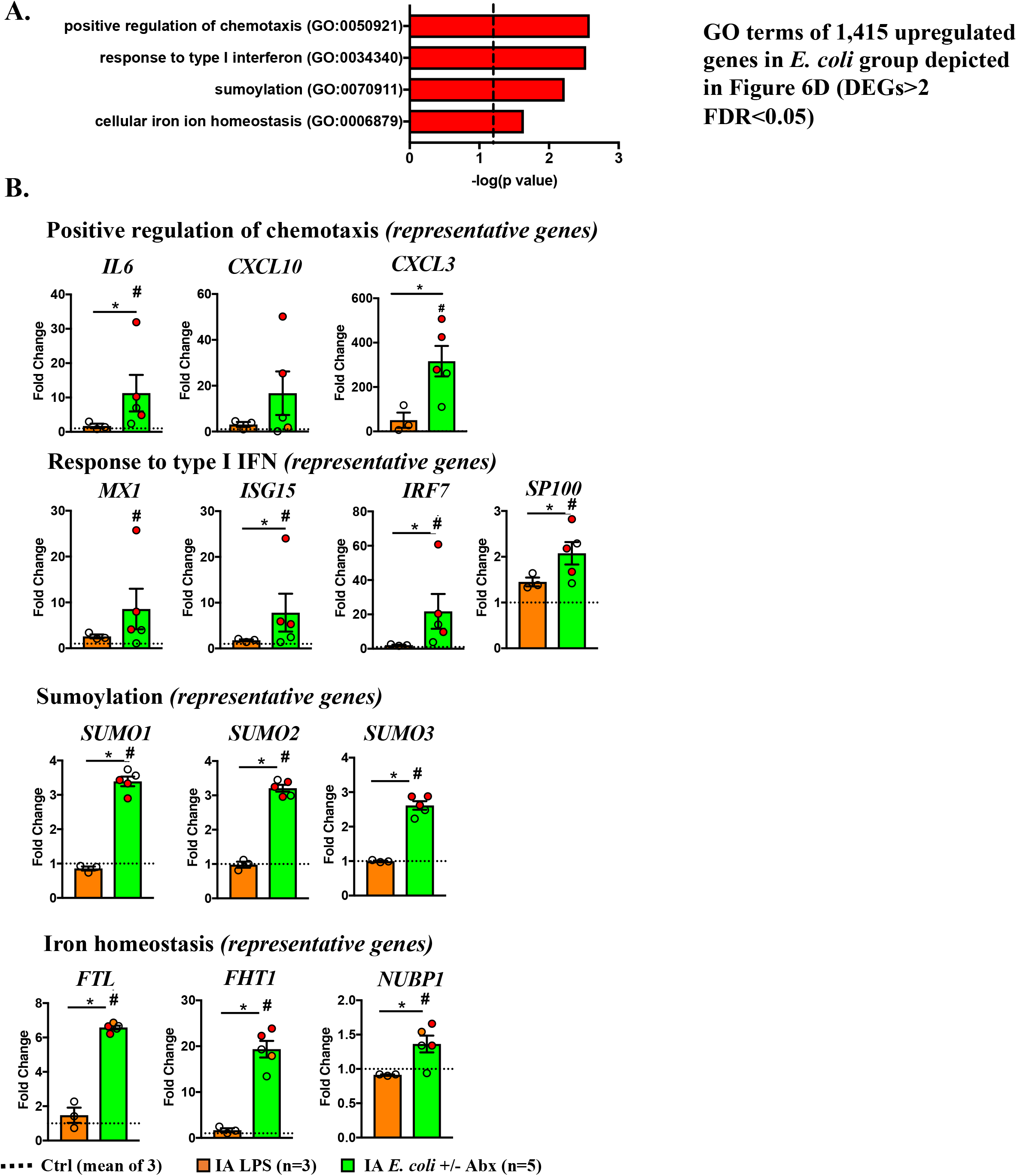
Upregulated pathways (GO terms) in chorioamnion-decidua by live *E. coli*. (**A**) Biological processes significantly upregulated in chorioamnion-decidua of *E. coli* treated animals compared to LPS exposure. (**B**) Representative genes of the biological processes shown in (A). Sequencing depth normalized counts from LPS and *E. coli* groups are expressed as fold change compared to the control group (dashed line representing the mean value of 3 ctrl animals). Red circles denote animals with PTL. Clear circles denote animals without PTL. *p<0.05 LPS vs. *E. coli*; #p<0.05 vs. control (Mann–Whitney U-test).

### Pathways for preterm labor induction

Within the *E. coli* exposed group, only 2/8 animals did not undergo PTL (both received antibiotics). These 2 samples gave us the opportunity to extend our transcriptomic analyses, by restricting the comparison between RNA-seq analyses from chorioamnion-decidua of animals infected by *E. coli* with or without PTL. Despite the paucity of samples, our data of PCA of global gene-expression profiles clearly demonstrated distinct clustering and heat maps of PTL vs. no PTL cases (**Supplementary Figure 6A-B**). We found 1,331 upregulated genes in PTL compared to no PTL samples (**Supplementary Figure 6C**). These genes were enriched in pathways such as inflammatory response, regulation of IL-6, IL-1 and TNF and other cytokine production, neutrophil mediated immunity and degranulation, NF-kB signaling and import to the nucleus, response to type I interferon and MyD88-indipendent toll-like signaling pathway (**Supplementary Figure 7A**). Downregulated genes in the PTL samples were enriched in cellular response (**Supplementary Figure 7B**), Notch signaling and T differentiation while mRNA-related processing and post-translational modification remained unaltered (**Supplementary Figure 7C**).

## Discussion

Viviparous species have evolved a strategy of inflammation inducing labor (22, 23). A corollary of the inflammation hypothesis is that there is a threshold of inflammation that triggers labor (14). Whether these concepts extend to inflammation-mediated preterm labor are not known. The peculiar anatomy of the female genital tract lends itself to vulnerability for ascension of the lower genital organisms in the upper genital space. An exuberant immune response to invading pathogens can incur collateral tissue injury and preterm labor (24). Therefore, the host immune system must balance the risk of inflammation induced preterm labor with protection from infectious organism. To gain insights into the pathogenesis of preterm labor, we used nonhuman primate models of IUI with and without PTL simulating human chorioamnionitis. We suggest that both the magnitude of inflammation and activation of certain pathways (e.g. IL-6, CCL2, prostaglandins) play a key role in triggering preterm labor (**Figure 8**).

**Figure 8.**
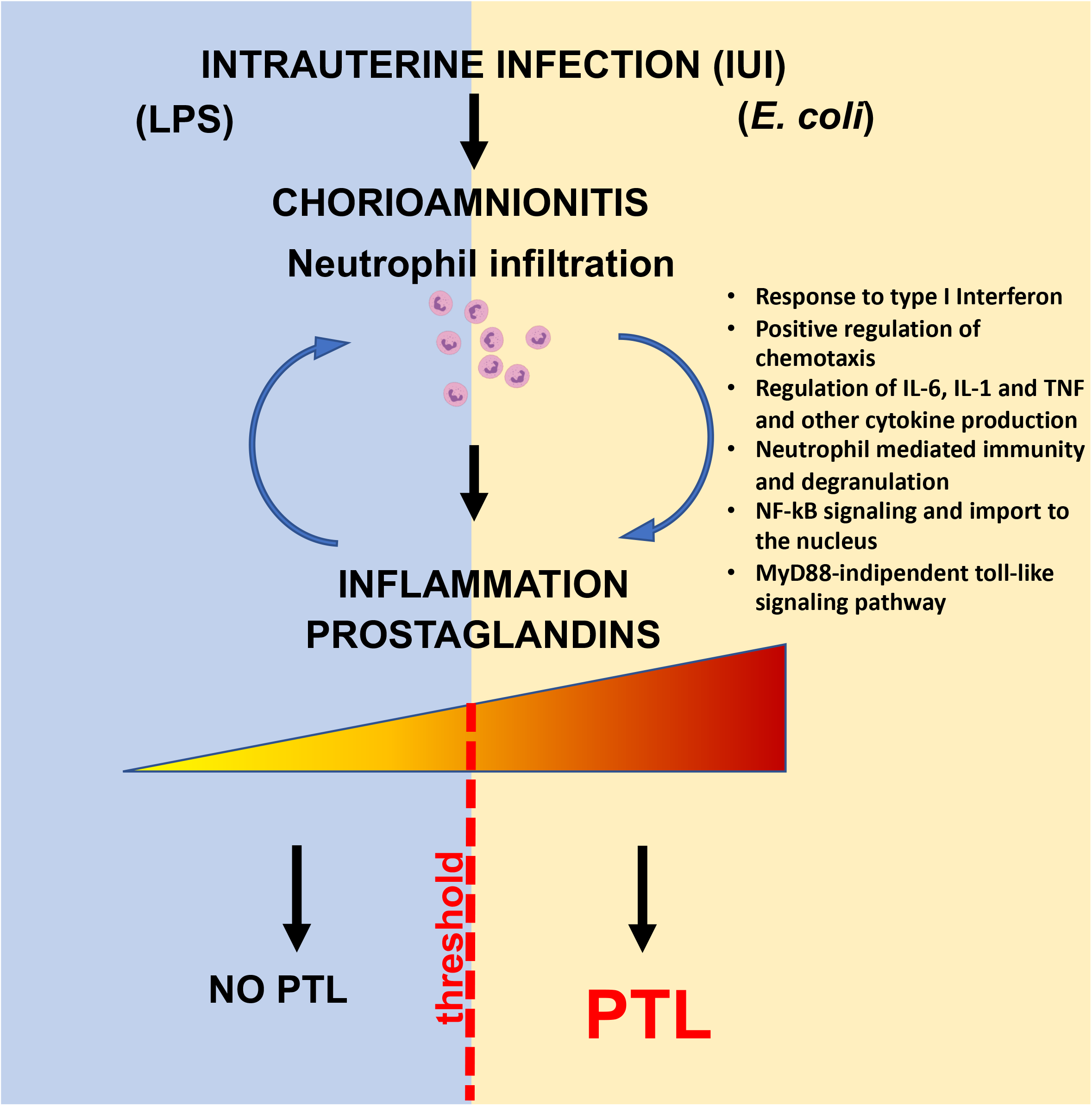
Working model: Intrauterine Infection and inflammation (IUI) are major risk factors for PTL. Chorioamnionitis is the hallmark feature of IUI and is typically characterized by the infiltration of neutrophils into the fetal membranes. Evidence suggests that the extent of bacterial colonization, route of infection, and the stimulatory capacity of the bacteria all play key roles in the activation of pro-inflammatory signaling cascades which induce production of pro-inflammatory cytokines (e.g., IL-6) and chemokines (e.g., CCL2). The intensity of immune response is a key player for the outcome of the pregnancy. which in turn promote prostaglandin production leading to preterm labor (PTL).

Although a number of pro-inflammatory cytokines are upregulated both by LPS and live *E. coli*, IL-6 and prostaglandins levels increased significantly more after *E. coli*. Mice with IL6 genetic knock out have delayed increases in prostaglandins, delayed onset of normal parturition despite on time progesterone withdrawal, and resistance to LPS induced preterm delivery (25, 26). In humans, higher amniotic fluid IL-6 predicted earlier preterm delivery (3, 27). However, exogenous administration of IL-6 alone, did not induce preterm labor in either mice or Rhesus macaques (28, 29). Thus, IL-6 signaling in the context of an inflammatory response seems to be required to trigger PTL. A number of different knock outs for genes in the prostaglandin signaling pathway in mice demonstrates the critical need for prostaglandin signaling in lysis of the corpus luteum of the ovary leading to progesterone withdrawal and normal parturition in mice (30). In humans, prostaglandin levels increase prior to parturition and administration of prostaglandins can induce preterm labor (31, 32). Thus, IL-6 and prostaglandin differentially upregulated by *E. coli* being potential factors for causing preterm labor is strongly supported by biologic rationale.

Differential responses to inflammatory stimuli may include different cell/tissues participating and/or different quantitative or qualitative host response. We observed that neutrophil infiltration in the amnion and *IL6* expression in the amnion was present after live *E. coli* stimulation but not after LPS injection in the amniotic fluid. Histological demonstration of neutrophils in the amnion denotes a higher stage of chorioamnionitis in humans compared to neutrophils only in the chorio-decidua (33). Although amnion cells can express *PTGS2* during normal labor (34), the source of prostaglandins during inflammation mediated preterm labor is not well understood. Both in our LPS and live *E. coli* models, we demonstrated that activated neutrophils expressing MPO were the major cells expressing inducible PTGS2 expression in the fetal membranes. Interestingly, *IL6* expression in the amnion was detected after live *E. coli* but not after LPS exposure. The cellular source of *IL6* was amnion epithelial cells and amnion mesenchymal cells with no contribution from the infiltrating neutrophils.

The quality and quantity of signals that trigger different inflammatory response at the maternal fetal interface are not well explored. Innate immune cells are known to modulate their responses based on input stimuli. As an example, the dynamics of NFkB activation in macrophages depends on which pattern recognition receptor (PRR) signaling is triggered and whether a combination of PRRs are signaled (35). Consistently, in our study, live *E. coli* with a combination of different PRRs induced more NFkB activation in the fetal membranes compared to LPS alone. Another mechanism that might explain a higher inflammatory response with live organism is that bacterial mRNA is known to induce a more potent innate immune response (36). Effector cells in the host can mount a tiered inflammatory response by switching on or off specific modules of transcription factors in response to different signals (37). Important classes of transcription factor differentially regulated by live *E. coli* include regulation of chemotaxis, type I IFN axis, sumoylation and iron homeostasis (**Figure 7**).

Using genetic knock out mice and antibody neutralization, we previously demonstrated that Type IFN axis primes LPS responses on maternal hematopoetic cells, increase the expression of *IL6* and *TNF* and increase susceptibility to preterm birth (26). Furthermore, we observed that type I interferon priming of LPS responses are conserved in non-human primates and humans (26). Type I interferon signaling can also increase the chemoattractant CXCL10-driven neutrophil recruitment to the site of inflammation (38). Thus, the higher type I interferon response in live *E. coli* may be an important driver of PTL in our study.

Sumoylation is a post-translational modification by small ubiquitin like modifiers (SUMO) proteins, that regulates innate immune response predominantly by negative regulation of IRF, type I interferon responses and inflammation (39–41). Our data suggest that *E. coli* increased the SUMO gene expression in the fetal membranes, which would be expected to decrease inflammation. This is counter to the increased inflammation observed in our study. However, SUMO expression is increased in the placenta in pre-eclampsia and chorioamnionitis (42, 43). Since sumoylation regulates function of a vast array of proteins, more work is needed to precisely understand how SUMO genes regulate inflammation during preterm labor.

Inflammation has a potent effect on iron homeostasis. Known as hypoferremia of inflammation, the cytokine-driven increase in hepcidin, largely mediated by IL6, decreases iron transport into plasma (44). The net effect is to decrease availability of nontransferrin bound free iron, which stimulates the growth of certain pathogenic bacteria. We recently demonstrated that in a Rhesus macaque model of IUI-induced by LPS, the fetus responded by rapidly upregulating hepcidin and lowering iron in fetal blood, without altering amniotic fluid iron status (45). The present study demonstrates a differential regulation of genes involved in iron homeostasis after live *E. coli* exposure. Overall, the findings suggest that the effects of iron homeostasis are likely due to bacterial infection from an invasive organism and innate host defense response.

A relatively large number of genes are differentially regulated (both up and down) in the chorioamnion-decidua of LPS vs. live *E. coli* exposed Rhesus macaques. However, in both screens (LPS vs. live *E. coli* and animals undergoing PTL vs no PTL within the *E. coli* exposed group), only a handful of differentially regulated pathways were represented. In both screens, IL6, NFkB, type I interferon, and neutrophil mediated immunity genes were represented. In sum, our findings potentially provide possible explanations for the role of both qualitative and quantitative thresholds of inflammation in induction of preterm labor. Our study provides further insights into the pathogenesis of intrauterine inflammation driven PTL.

## MATERIALS AND METHODS

### Animals

Normally cycling, adult female rhesus macaques (*Macaca mulatta*) were time mated. At ~130 d of gestation (~80% of term gestation), the pregnant rhesus received either 1 ml saline solution (n=12) or 1 mg LPS (Sigma-Aldrich, St. Louis, MO in 1 ml saline solution (n= 8) or live uropathogenic *E. coli* derived from a cystitis clinical isolate (strain UTI89) (10^6^ CFU in 1 mL) (n=5). All injections were given by ultrasound-guided intraamniotic (IA) injection. In a subset of *E. coli* injected animals, antibiotics were given starting 24h after bacterial injection (n=8). The antibiotic dosing regimen was: Intramuscular (IM) cefazolin 25 mg/k twice daily + IM enrofloxacin 5mg/k twice daily. Additionally, IA injection of cefazolin (10mg) + IA enrofloxacin (1mg) were given once daily. Dams injected with LPS were surgically delivered 48h later while dams infected with *E. coli* had preterm labor generally about 2d after IA *E. coli* injection. The animals given antibiotics delivered preterm 2-3d after *E. coli* injection (n=6) with the remaining two animals delivered surgically at 3d without preterm labor. Only 1 animal in the *E. coli* group delivered vaginally and the remainder 4 animals were surgically delivered in active preterm labor. After delivery, fetuses were euthanized with pentobarbital, and fetal tissues were collected. Some control and LPS animals were used in a previous study (21). The animals used for each experiment are listed in **Supplementary Table 1**.

### Chorioamnion-decidua dissection

Extra-placental membranes were dissected away from the placenta as previously described (17, 21). After scraping decidua parietalis cells with the attached chorion, the amnion tissue was peeled away from the chorion with forceps. Chorio-decidua cells were washed, and digested with Dispase II (Life Technologies, Grand Island, NY) plus collagenase A (Roche, Indianapolis, IN) for 30 min followed by DNase I (Roche) treatment for another 30 min. Cell suspensions were filtered, the red blood cells lysed and prepared for flow cytometry or FACS-sorting. Viability was >90% by trypan blue exclusion test. Tissues of fetal membranes were used for RNA analyses described below.

### Flow cytometry of chorio-decidua cells

Monoclonal antibodies (mAbs) used for multiparameter flow cytometry (LSR Fortessa 2, BD Biosciences, San Diego, CA) and gating strategy to identify the different leukocyte subpopulations was done as previously described (21) (and **Supplementary Figure 2**). Cells were treated with 20 μg/mL human immunoglobulin G (IgG) to block Fc receptors, stained for surface markers for 30 min at 4°C in PBS, washed, and fixed in fixative stabilizing buffer (BD Bioscience). Samples were acquired within 30 minutes after the staining. All antibodies were titrated for optimal detection of positive populations and mean fluorescence intensity. At least 500,000 events were recorded for each sample. Doublets were excluded based on forward scatter properties, and dead cells were excluded using LIVE/DEAD Fixable Aqua dead cell stain (Life Technologies). Unstained and negative biological population were used to determine positive staining for each marker. Data were analyzed using FlowJo version 9.5.2 software (TreeStar Inc., Ashland, OR).

### Chorioamnion-decidua tissue cytokine Quantitative RT-PCR

Total RNA was extracted from snap-frozen chorioamnion-decidua and amnion biopsies after homogenizing in TRIzol (Invitrogen). RNA concentration and quality were measured by Nanodrop spectrophotometer (Thermo-Scientific). Reverse transcription of the RNA and quantitative RT-PCR were performed using qScript One-Step RT-qPCR Kit (Quanta Biosciences), following the manufacturer’s instructions and with Rhesus-specific TaqMan gene expression primers (Life Technologies). Eukaryotic 18S rRNA (Life Technologies) was endogenous control for normalization of the target RNAs.

### RNA sequencing and analysis

Total RNA was purified, treated with DNase using RNeasy mini kit following manufacturer’s recommendation (Qiagen, Valencia, CA, USA). After purification, the concentration of total RNA was measured using Nanodrop ND-1000 spectrophotometer (Thermo Fisher Scientific, Wilmington, DE, USA) and the quality was analyzed by the Bioanalyzer 2100 (Agilent Technologies, Santa Clara, CA, USA). Samples with RNA Integrity Number (RIN) ≥ 9.0 were used for mRNA sequencing. RNA-library preparation was performed at DNA core facility at Cincinnati Children’s Hospital Medical Center. Single-end read sequencing by Illumina HiSeq2500 Ultra-High-throughput sequencing system (Illumina Inc. San Diego, CA, USA) was used at an average depth of 20 million reads per sample. Raw sequences were accepted once they passed the quality filtering parameters used in the Illumina GA Pipeline.

The reads were mapped with STAR 2.5.3a to the *Macaca mulatta* genome (Mmul 8.0.1). The counts for each gene were obtained using quantMode GeneCounts in the STAR commands, and only counts for the featured genes were reserved. Differential expression analyses were carried out using DESeq2. PCA analysis were performed on the rlog counts from DESeq2 using the R function prcomp. Volcano plots were made for the results from the differential expression analyses. Heatmaps were plotted on the log2 value of the normalized counts. Inference of biological processes were generated using Enrichr (46).

### IL6, PTGS2 mRNAs and MPO staining

RNAscope^®^ (RNAscope^®^ Fluorescent Multiplexed reagent kit, Advanced Cell Diagnostics, United States) was used as per manufacturer’s protocol and adjusted for dual detection of mRNA and protein. RNAscope Multiplex Fluorescent Reagent Kit V2 (323100, RNAscope^®^, Advanced Cell Diagnostics, Inc) was used according to manufacturer’s protocol to label mRNAs for *IL6* and *PTGS2*. Five μm paraffin embedded sections of the chorion-amnion decidua tissue were stained with positive controls (300040, Advanced Cell Diagnostics, Inc), negative controls (320871, Advanced Cell Diagnostics, Inc), and probes for targeting Rhesus PTGS2 (497758-C2, Advanced Cell Diagnostics, Inc) and Rhesus IL6 (310378, Advanced Cell Diagnostics, Inc). Probes were fluorescently labeled with Opal™ 570 Reagent (Akoya Biosciences, FP1488001KT) and Opal™ 620 Reagent (Akoya Biosciences, FP1487001KT) and stained with DAPI (4’,6-diamidino-2-phenylindole) to demarcate the nucleus. Following completion of the RNAscope^®^ protocol, sections were immunolabelled using a rabbit anti-MPO polyclonal affinity purified antibody (1:500; Dako Omnis, A0398) and incubated with Dako anti-rabbit HRP polymer (Agilent, K4003) and then conjugated with Opal™ 690 Reagent (Akoya Biosciences, FP1497001KT). Then slides were digitalized on a Leica Aperio Versa (Leica Biosystems, Inc., Vista, CA).

### Cytokines and Prostaglandins ELISA

Cytokine/chemokine concentrations in AF, fetal, and maternal plasma were determined by Luminex using non-human primate multiplex kits (Millipore). Lipids were extracted from the AF using methanol to measure prostaglandins PGE2 (Oxford Biomedical Research, Oxford, MI) and PGF2α (Cayman Chemical, Ann Arbor, MI).

### Endotoxin assay

Endotoxin level in the AF was determined by Limulus amebocyte lysate assay (LAL; Lonza) according to the test procedure recommended by the manufacturer’s instructions.

### Histology of fetal membranes for chorioamnionitis

H&E staining for both Rhesus and human fetal membrane was performed, and pictures were taken. H&E-stained sections of human fetal membranes were scored in a blinded manner (by SGK) for chorioamnionitis using criteria outlined by Redline et al (47).

### Statistics

Prism version 7 software (GraphPad) was used to analyze data. Values were expressed as means ± SEM. Two-tailed Mann-Whitney U tests (for nonnormally distributed continuous variables) and Fisher’s exact test for categorical variables were used to determine differences between groups. Results were considered significant for p ≤ 0.05.

### Study approval

All animal procedures were approved by the IACUC at the University of California Davis.

## ACKNOWLEDGEMENTS

This study was supported by U01 ES029234 (CAC), Burroughs Wellcome grant (CAC), CCHMC Perinatal Infection and Inflammation Collaborative (CAC), R21HD90856 (SGK), and R01HD 98389 (SGK). We thank Sarah Davis, Jennifer Kendrick, Sarah Lockwood, Anne Gibbons, Paul-Michael Sosa, and Marie Jose-Lemoy research personnel at the CNPRC, U.C. Davis for help with the animals.

## AUTHOR CONTRIBUTIONS

MC, PP, PS, FM, LAM, MP, AHJ, SD, SSW, CAC, and SGK participated in data generation. MC, PP, PS, FM, MP, AHJ, CAC, and SGK participated in analysis and interpretation of data. AHJ, CAC, and SGK participated in the conception and design of the study and obtained the funding. MC and SGK wrote the manuscript. All authors have reviewed the manuscript and approve the final version.

---

**Supplementary Table 1.**
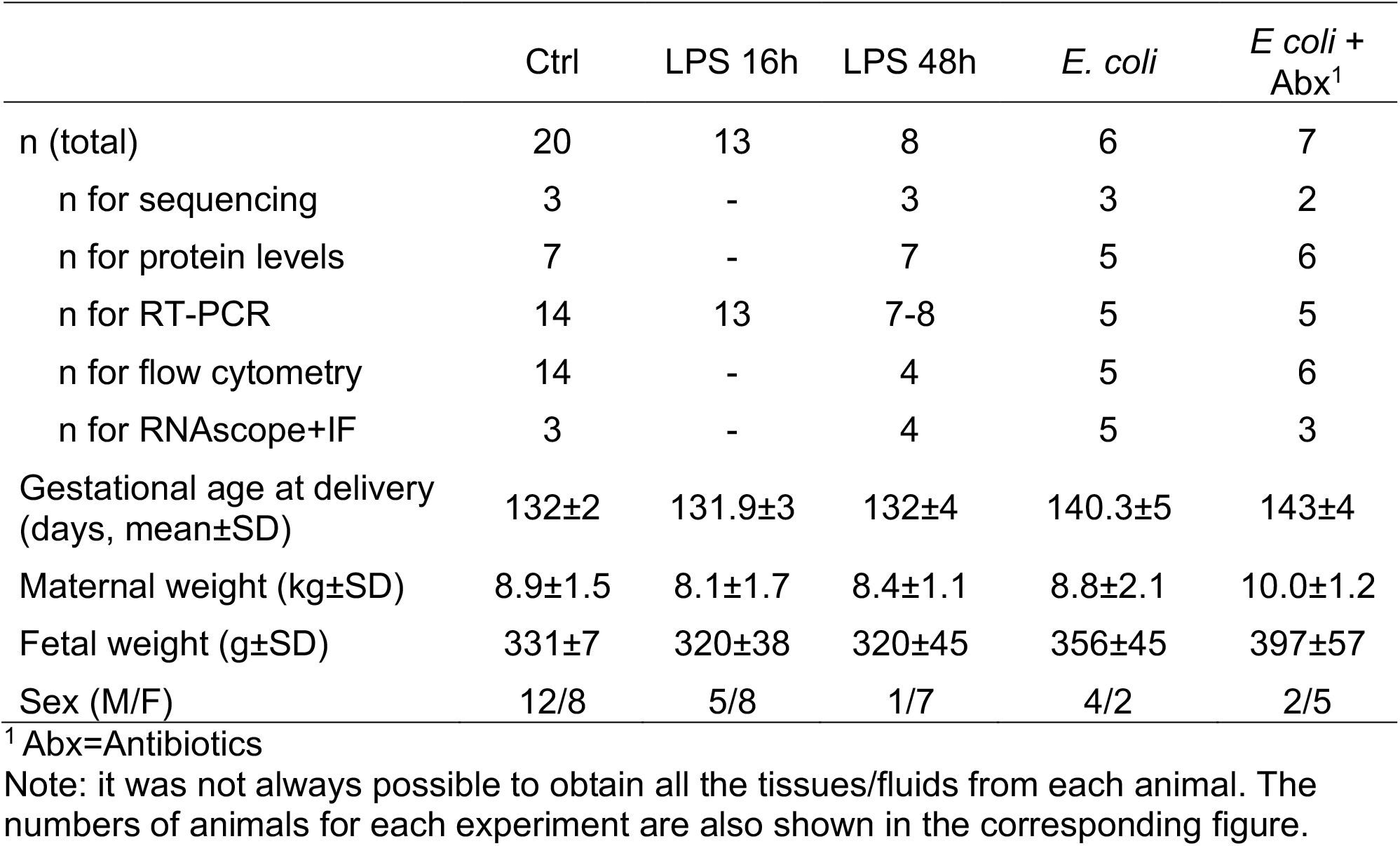
Animal data and sample sizes

|  | Ctrl | LPS 16h | LPS 48h | *E. coli* | *E. coli* +<br>Abx<sup>1</sup> |
| --- | --- | --- | --- | --- | --- |
| n (total) | 20 | 13 | 8 | 6 | 7 |
| n for sequencing | 3 | - | 3 | 3 | 2 |
| n for protein levels | 7 | - | 7 | 5 | 6 |
| n for RT-PCR | 14 | 13 | 7-8 | 5 | 5 |
| n for flow cytometry | 14 | - | 4 | 5 | 6 |
| n for RNAscope+IF | 3 | - | 4 | 5 | 3 |
| Gestational age at delivery<br>(days, mean±SD) | 132±2 | 131.9±3 | 132±4 | 140.3±5 | 143±4 |
| Maternal weight (kg±SD) | 8.9±1.5 | 8.1±1.7 | 8.4±1.1 | 8.8±2.1 | 10.0±1.2 |
| Fetal weight (g±SD) | 331±7 | 320±38 | 320±45 | 356±45 | 397±57 |
| Sex (M/F) | 12/8 | 5/8 | 1/7 | 4/2 | 2/5 |

<sup>1</sup> Abx=Antibiotics
Note: it was not always possible to obtain all the tissues/fluids from each animal. The numbers of animals for each experiment are also shown in the corresponding figure.

**Supplementary Figure 1.**
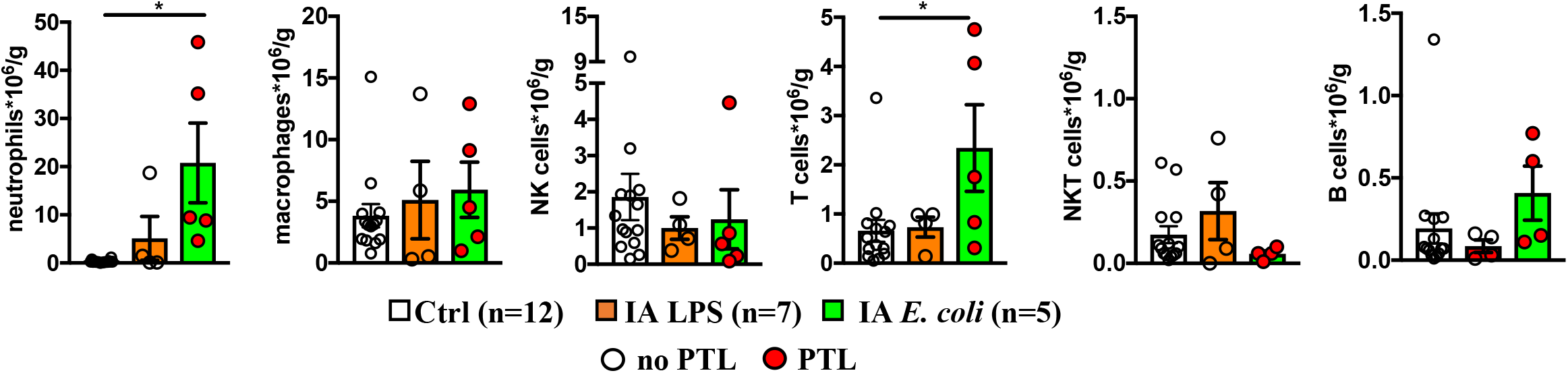
Chorio-decidua immune cells upon different IA exposures. Chorio-decidua cell suspensions were analyzed by multiparameter flow cytometry. Cell count is expressed per gram of tissue. Data are mean, SEM, *p<0.05 between comparators (Mann-Whitney U-test). Red circles denote animals with PTL. Clear circles denote animals without PTL.

**Supplementary Figure 2.**
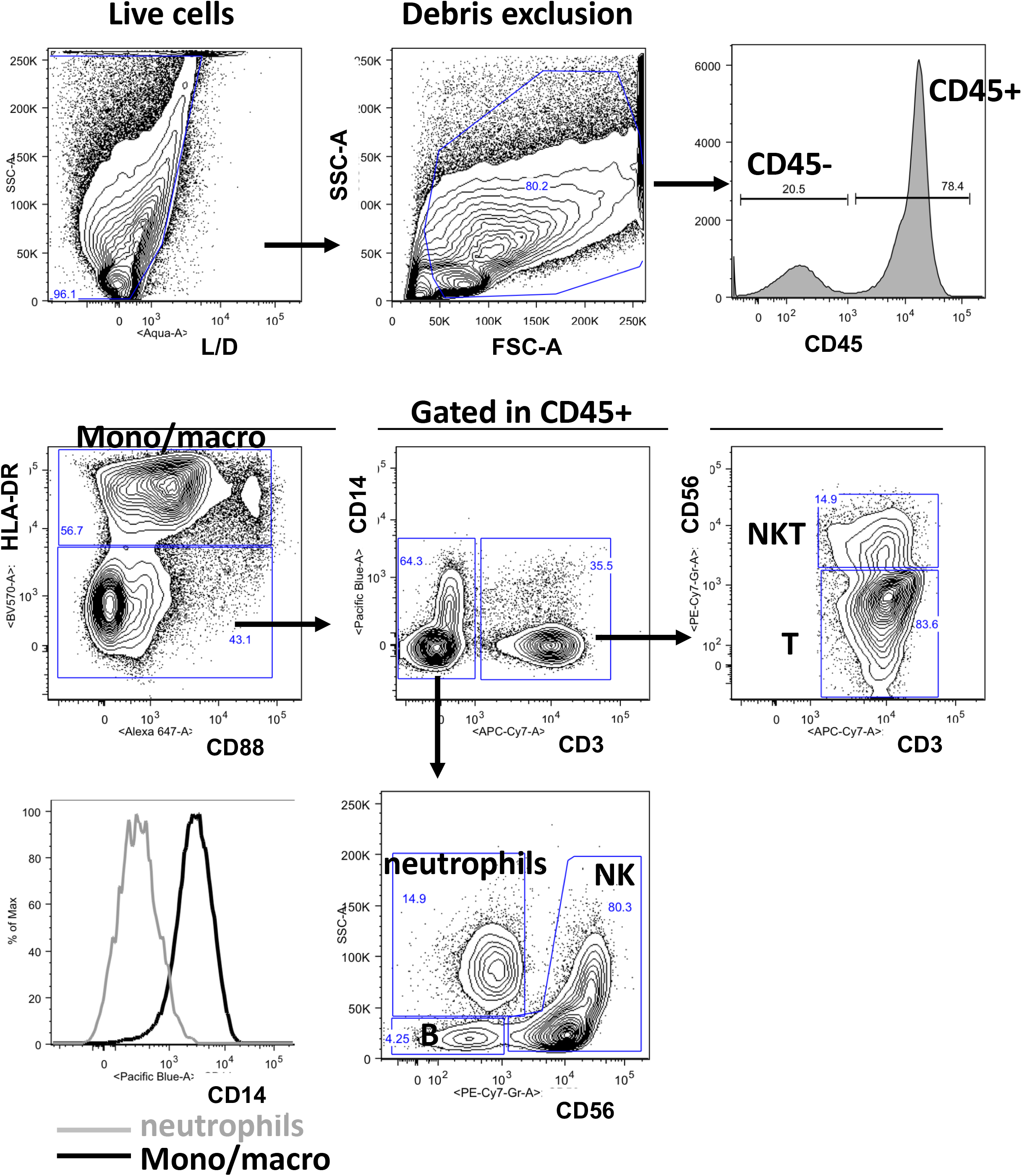
Flow cytometry phenotype of chorio-decidua cells. Chorio-decidua parietalis cells were scraped, digested with protease/DNAase and single cell suspensions were used for multi-parameter flow cytometry analysis. Briefly, live cells were first identified by the absence of LIVE/DEAD stain and forward-/side-scatter expression, excluding cell debris. Leukocytes were gated as CD45+ cells.

**Supplementary Figure 3.**
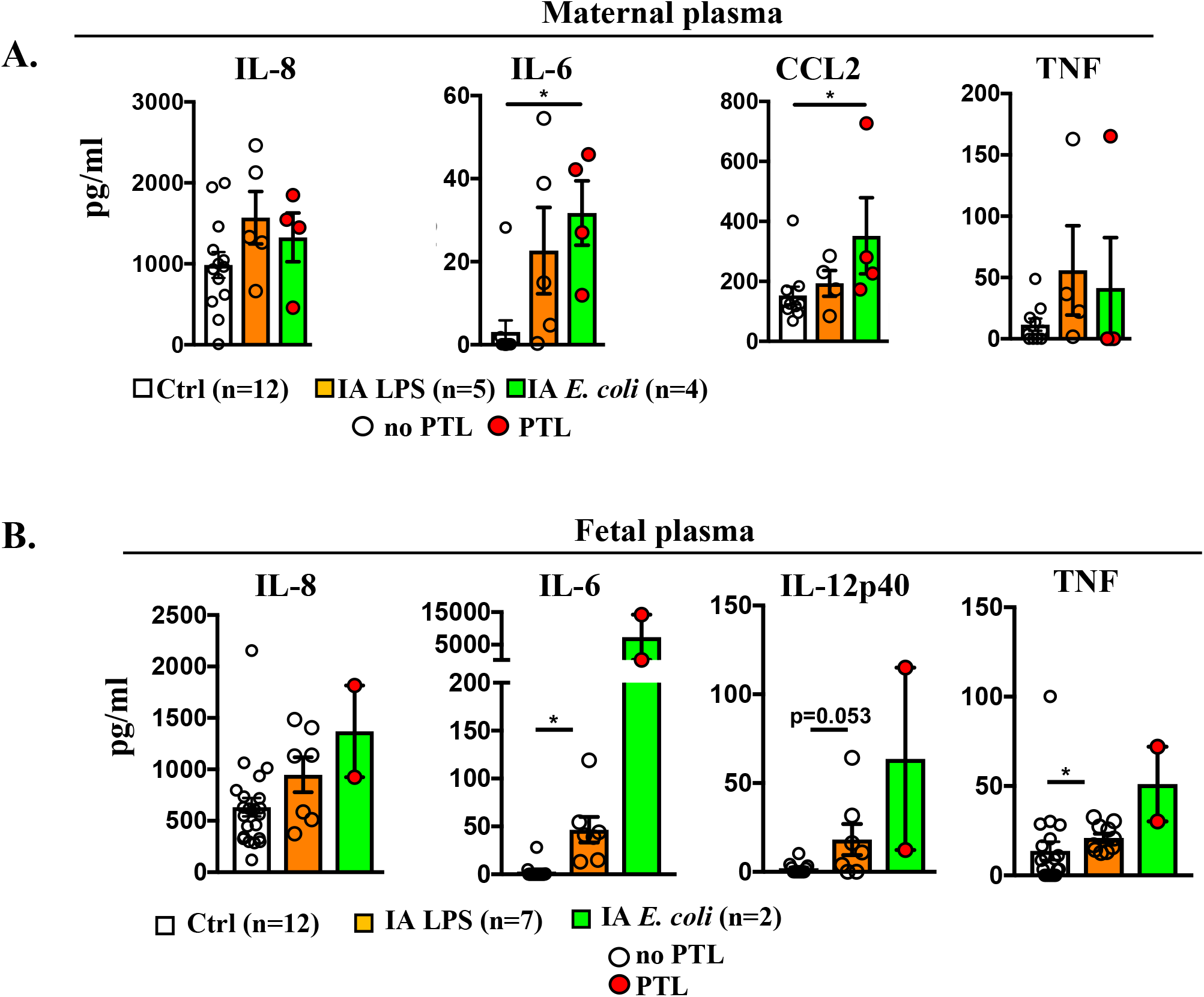
Inflammatory cytokines in maternal and fetal plasma. (**A**) Maternal plasma and (**B**) Fetal cord blood inflammatory cytokine in different Rhesus IUI models were measured by multiplex Elisa. Compared to control values, only CCL2 modestly increased in the the maternal plasma of IA *E. coli* group. In the fetal plasma, IL-6 and IL12p40 increase in both IA *E.coli* and IA LPS groups compared to saline controls. Data are mean, SEM, *p<0.05 between comparators (Mann-Whitney U-test). Red circles denote animals with PTL. Clear circles denote animals without PTL.

**Supplementary Figure 4.**
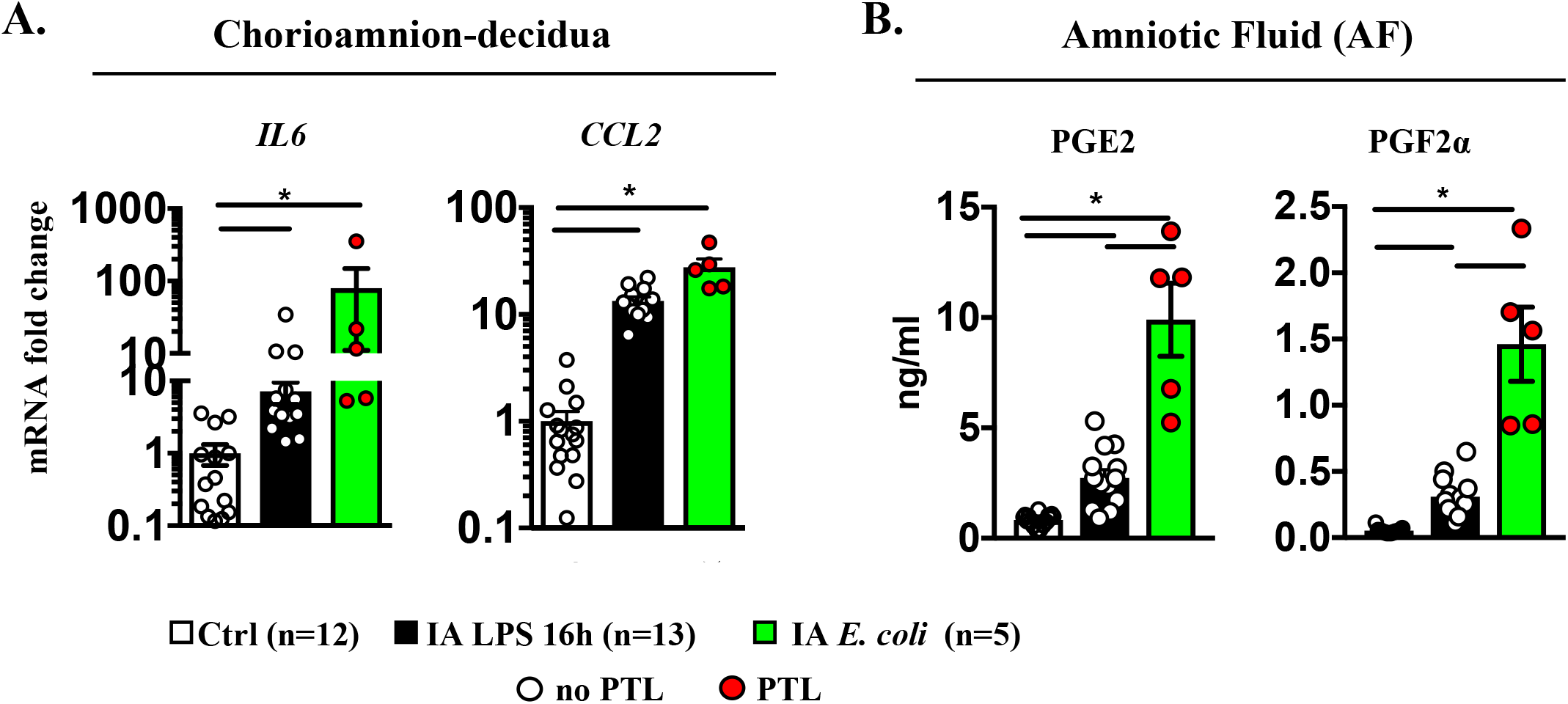
Higher IL-6 and CCL2 mRNA expression and Prostaglandins by live *E. coli* compared to LPS 16h exposure. Rhesus were treated as in Fig. 1A but LPS exposure was for 16h prior to surgical delivery. (**A**) Chorio-amnion decidua inflammatory cytokine mRNAs in different Rhesus IUI models were measured by qPCR. Average mRNA values are fold increases over the average value for control after internally normalizing to the housekeeping 18S RNA. (**B**) Prostaglandin levels in different Rhesus IUI models were measured by multiplex ELISA in the amniotic fluid. Data are mean, SEM, *p<0.05 between comparators. Red circles denote animals with PTL. Clear circles denote animals without PTL.

**Supplementary Figure 5.**
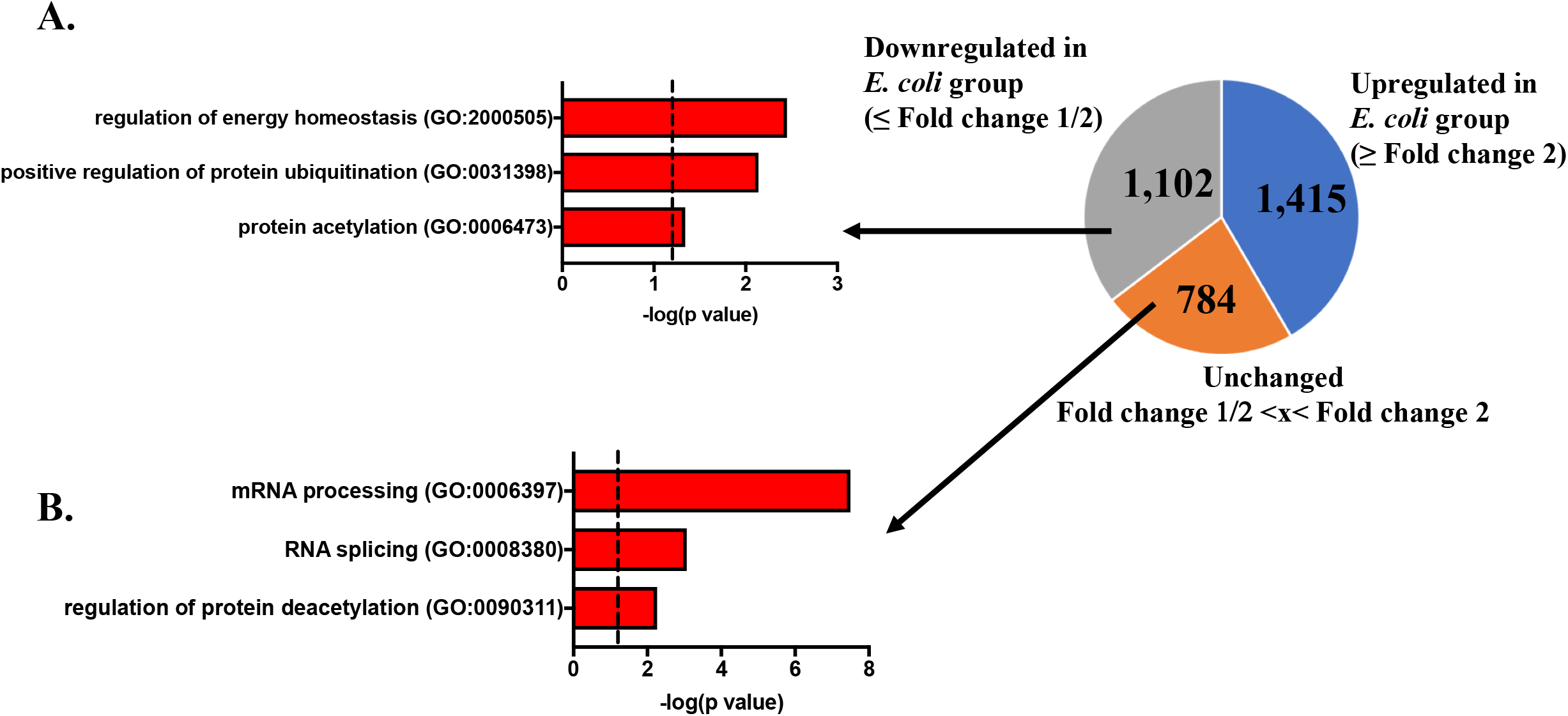
Downregulated and unchanged GO terms in chorioamnion-decidua by live *E. coli*. (**A**) Biological processes significantly downregulated in chorioamnion-decidua of *E. coli* treated animals (DEGs ≤ Fold change 1/2) or (**B**) unchanged (Fold change 1/2 <x< Fold change 2) vs. LPS.

**Supplementary Figure 6.**
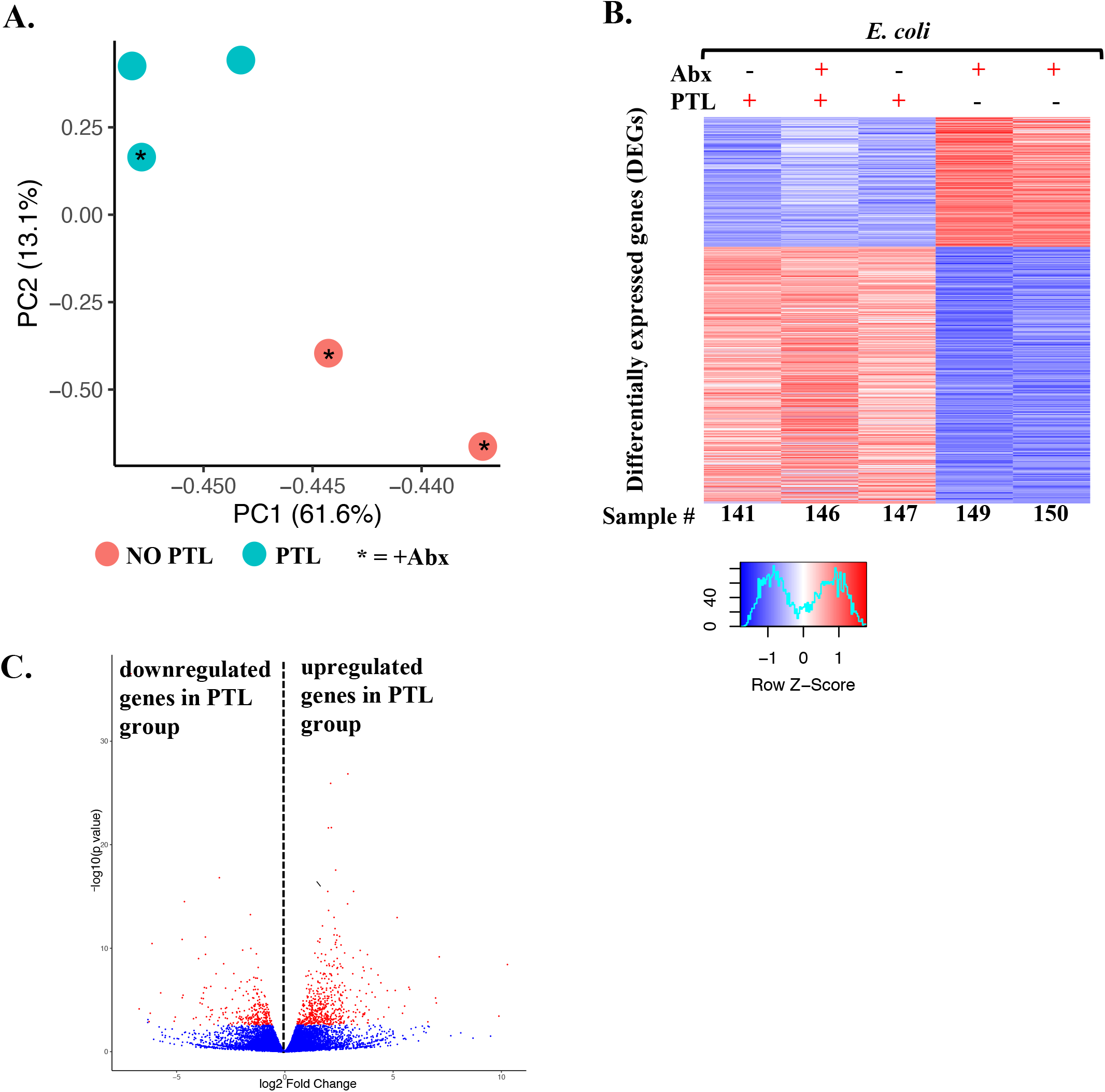
Comparative transcriptomics in chorioamnion-decidua between IA *E. coli-exposure* with or without PTL. (**A**) PCA of RNA-seq data from chorioamnion-decidua cells showing different clustering based on the presence of PTL. (**B**) Heatmap of genes that are differentially expressed (DEGs) in chorioamnion-decidua cells of animals with PTL vs. NO PTL. (**C**) Volcano plot displaying genes detected by RNA-seq in chorioamnion-decidua from *E. coli* treated animals with PTL compared to NO PTL animals.

**Supplementary Figure 7.**
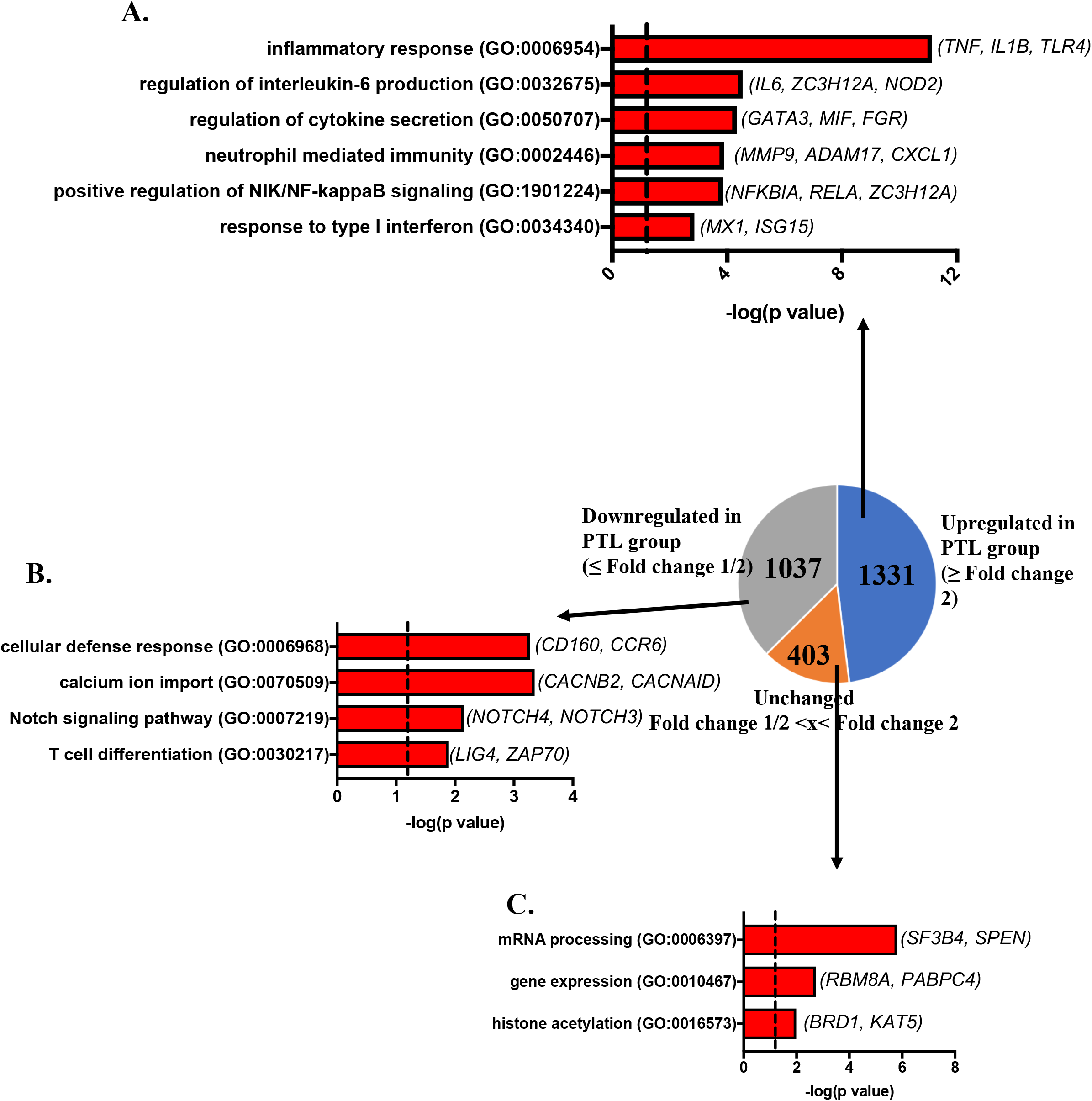
Animals undergoing PTL show upregulated pathways leading to increased inflammatory response. Pie chart displaying DEGs upregulated (Fold Change ≥ 2), unchanged (Fold change 1/2 <x< Fold change 2) and downregulated in PTL group (Fold Change ≤ 1/2). Gene ontology pathways and representative genes in (**A**) upregulated in animals with PTL, (**B**) downregulated (DEGs ≤ Fold change 1/2) or (**C**) unchanged in *E. coli* treated animals with vs. without PTL.

